# Connecting species-specific extents of genome reduction in mitochondria and plastids

**DOI:** 10.1101/2023.12.14.571654

**Authors:** Konstantinos Giannakis, Luke Richards, Kazeem A. Dauda, Iain G. Johnston

## Abstract

Mitochondria and plastids have both dramatically reduced their genomes since the endosymbiotic events that created them. The similarities and differences in the evolution of the two organelle genome types has been the target of discussion and investigation for decades. Ongoing work has suggested that similar mechanisms may modulate the reductive evolution of the two organelles in a given species, but quantitative data and statistical analyses exploring this picture remain limited outside of some specific cases like parasitism. Here, we use cross-eukaryote organelle genome data to explore evidence for coevolution of mitochondrial and chloroplast genome reduction. Controlling for differences between clades and pseudoreplication due to relatedness, we find that mtDNA and ptDNA gene retention are related across taxa, in a generally positive correlation that appears to differ quantitatively across eukaryotes, for example, between algal and non-algal species. We find limited evidence for coevolution of specific mtDNA and ptDNA gene pairs, suggesting that the similarities between the two organelle types may be due mainly to independent responses to consistent evolutionary drivers.

## Introduction

Mitochondria and plastids (the class of organelles to which chloroplasts belong) are essential compartments in eukaryotic cells. Originally independent organisms, both organelle classes arose from endosymbiotic capture and subsequent evolution (Zimorski et al. 2014; Sagan 1967; Roger, Muñoz-Gómez, and Kamikawa 2017; Keeling 2010). This evolution has involved the dramatic reduction of the organelle genomes (mtDNA and ptDNA respectively; generally, organelle DNA or oDNA) (Roger, Muñoz-Gómez, and Kamikawa 2017; Keeling 2010; Hjort et al. 2010). This reduction has proceeded to different extents in different taxa (Giannakis et al. 2022; Johnston and Williams 2016; Janouškovec et al. 2017). Some jakobid protists retain over 60 protein-coding genes in their mtDNA (Lang et al. 1997) while some parasitic organisms retain only 3 and others have completely lost mtDNA (Hjort et al. 2010; Maciszewski and Karnkowska 2019; de Paula, Allen, and van der Giezen 2012). Some Rhodophyta contain around 200 protein-coding ptDNA genes (Janouškovec et al. 2013) while some parasitic plants contain only dozens (Naumann et al. 2016) and some reduced forms of plastids – like the apicoplasts found in Apicomplexans (McFadden 2011) – even fewer.

The diversity of organelle DNA gene profiles across eukaryotes raises, among others, two dual questions. First, what are the properties of a *gene* that make it more or less likely to be retained in a given species? And second, what are the properties of a *species* that make it more or less likely to retain a given gene? Numerous hypotheses for the first question have been proposed. It has been suggested that gene encoding hydrophobic products are preferentially retained in oDNA due to the difficulty of importing remotely-encoded hydrophobic proteins into the organelle (von Heijne 1986; Björkholm et al. 2015), including via mistargeting of mitochondrial products expressed in the cytosol – demonstrated across different genes (Björkholm et al. 2017). The colocation for redox regulation (CoRR) hypothesis proposes that retaining essential genes in organelles – where they can be expressed in response to local stimuli – supports local control of redox poise and organelle behaviour rather than relying on slow and indirect nuclear transport (Allen 2015; Allen and Martin 2016; de Paula, Allen, and van der Giezen 2012). Economic considerations of the energy required by different encoding locations also support organelle retention of some genes (Kelly 2021). A role for organelle genes as long-term redox sensors, identifying bioenergetically competent cells, has been proposed (Wright, Murphy, and Turnbull 2009). Data-driven comparison of hypotheses using cross-eukaryotic genome data has provided support for the retention of genes encoding hydrophobic products, and products which are physically central in their functional complexes (supporting CoRR-like control) (Giannakis et al. 2022; Johnston and Williams 2016). Pronounced convergence in oDNA ribosomal protein-coding gene content across species has been found in support of CoRR-like pressures on gene retention (Maier et al. 2013).

Some of these mechanisms – likely in combination (Giannakis et al. 2022; Johnston and Williams 2016) – may explain some of the gene-to-gene variability in retention. But what of the dual species-to-species variability in number of oDNA genes retained? It is well known that many parasites atrophy organelle genomes to very reduced or even absent oDNA, likely due to a reduced energetic demand resulting from their exploitation of host metabolism (Keeling 2010; Hjort et al. 2010; Roger, Muñoz-Gómez, and Kamikawa 2017; Giannakis et al. 2022). Self-pollinating and clonal plant species typically transfer more oDNA genes to the nucleus; theoretical work has demonstrated that self-pollination accelerates such transfer when nuclear encoding is beneficial (Brandvain, Barker, and Wade 2007; Brandvain and Wade 2009). The coupled effects of nuclear genome reduction have been hypothesized as a reason for relative gene richness of red algal ptDNA (Qiu et al. 2017). The “limited transfer window” hypothesis places bounds on the time period during which a given taxa can transfer organelle genes (Barbrook, Howe, and Purton 2006).

Outside these specific cases, a general picture (if even applicable) has yet to fully emerge. One candidate is the “mutational hazard hypothesis” (Lynch, Koskella, and Schaack 2006), which ties gene retention to taxon-specific oDNA mutation rates, suggesting that organisms with low rates of oDNA mutation (for example, plants) can support more organelle-encoded content without risking deleterious effects of this content becoming mutated. This picture has support from some observations and competition from others (Smith 2016a), with several instances where the hypothesized relationship between mutation rate and coding content runs in a different way to that predicted. Another candidate contribution to a cross-species picture is an interpretation of CoRR which attempts to bridge intercellular and organismal scales. Here, when organelles are required to rapidly adapt to changing environmental conditions experienced by their “host”, they are predicted to retain more oDNA genes, to allow individual local control of organelles (Johnston 2019). Hence, organisms experiencing highly variable bioenergetic and metabolic demands from their environment would retain more oDNA genes, and those experiencing lower and more stable demands would retain fewer. This picture is supported to some extent by mathematical modelling (García-Pascual, Nordbotten, and Johnston 2022) and comparative analysis of links between ecological traits and oDNA gene counts (Giannakis, Richards, and Johnston 2023) – although we must point out that this picture does not and probably cannot explain the full diversity of oDNA retention profiles across eukaryotes. Although this theoretical picture predicts that similar factors shape – to some extent – mtDNA and ptDNA evolution, this prediction has to our knowledge yet to be tested with data. Here, while accepting that seeking too tight links between mitochondrial and plastid evolution can be misleading (Smith and Keeling 2015), we set out to explore whether a quantifiable relationship exists between mtDNA and ptDNA gene counts across eukaryotes.

## Methods

### Data gathering and curation

We used the pipeline from (Giannakis et al. 2022) to collect cross-eukaryote data on protein-coding oDNA genes from NCBI’s Organelle Genome database (O’Leary et al. 2016) and to curate and unify annotations. This pipeline involves resolving the inconsistent annotation of genes across eukaryotic records via two branches. First, an unsupervised all-against-all BLAST comparison (Camacho et al. 2009) of all oDNA CDS (coding sequence) records is performed, followed by iterative hierarchical clustering and relabelling based on the BLAST outputs, to give sets of genes that form a connected network in the space of BLAST comparison metrics (regardless of original annotation). Second, a supervised process of relabelling annotations based on a manually compiled dictionary of annotation protocols across taxa. The relabelled annotations from these two approaches are then compared and any inconsistencies manually resolved.

NCBI’s Common Taxonomy Tool (Federhen 2012) was used to estimate phylogenetic topology. The Encyclopedia of Life (Parr et al. 2014), Wikipedia (Wikipedia 2023), World Register of Marine Species (Ahyong et al. 2023), the USDA PLANTS Database (USDA and NRCS 2023), World Flora Online (WFO 2023), and AlgaeBase (Guiry and Guiry 2023) were used to collate ecological information on species.

### Statistical analysis and visualisation

We used *phytools* (Revell 2012), *ape* (Paradis and Schliep 2019), and *phangorn* (Schliep 2011) for phylogenetic construction and analysis. For phylogenetic linear models (PLM) we used the *phylolm* package (Tung Ho and Ané 2014) for linear regression accounting for phylogenetic correlation with generalized estimating equations (Paradis and Claude 2002). We used *nlme* (DebRoy 2006) and *lme4* (Bates et al. 2015) for linear mixed models and *pheatmap* (Kolde 2019) for clustering and analysis. We used *Oncotree* (Szabo and Boucher 2008) for inference of progression. *igraph* (Csardi and Nepusz 2006) provided network analysis and visualization support. For visualization we used *ggplot2* (Wickham 2011), *ggpubr* (Kassambara 2020), *ggrepel* (Slowikowski 2021), *ggraph* (Pedersen 2020), *ggtree* (Yu et al. 2017), *ggtreeExtra* (Xu et al. 2021), and *ggVennDiagram* (Gao, Yu, and Cai 2021).

## Results

### Profiles of mitochondrial and chloroplast gene count across eukaryotes

We used the pipeline from (Giannakis et al. 2022) to collect cross-eukaryote data on protein-coding oDNA genes from NCBI’s Organelle Genome database (O’Leary et al. 2016) and curate and unify annotations (see Methods). Following this pipeline, we found 205 unique species for which mitochondrial and plastid gene counts were available. 147 were Viridiplantae (green plants and algae), 30 Rhodophyta (red algae), 11 Bacillariophyta (diatoms), 9 Phaeophyceae (brown algae), 2 Apicomplexa (typically parasitic protists), and representatives of other algal groups: 2 Cryptophyceae, and 1 each of Bolidophyceae, Eustigmatophyceae, Glaucocystophyceae, and Haptista. A simple plot of mitochondrial (MT) gene count vs plastid (PT) gene count (Fig. 1A) shows considerable range and structure in the protein-coding gene count profiles. The Viridiplantae samples occupy a wide spread of MT counts at intermediate PT counts; Rhodophyta cluster at intermediate MT and very high PT counts; other algae occupy a high MT/high PT region.

**Figure 1.**
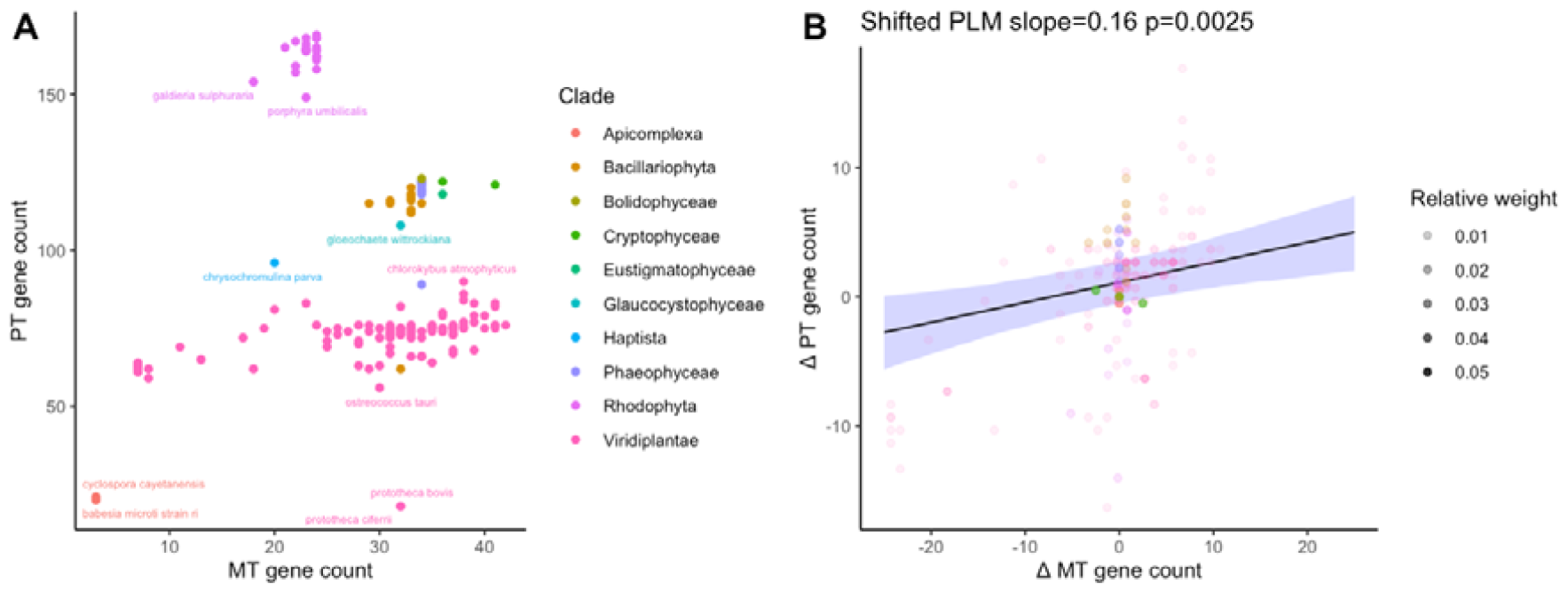
MT and PT gene counts across eukaryotes. (A) Raw protein-coding oDNA gene counts across species in our dataset, labelled by basal eukaryotic clade. Some species are labelled for illustration. (B) oDNA differences from clade average (ΔMT and ΔPT). Opacity of points in (B) is an illustration of their weighting in phylogenetic linear modelling (PLM): the inverse of the sum of their associated row in the variance-covariance matrix, reflecting a species’ relatedness to other members of the dataset. Sets of closely related species provide less independent information and so have a lower weighting. A PLM fit suggests a significant (p = 0.0025) relationship between ΔMT and ΔPT, also found in a linear mixed model for raw oDNA counts (Supplementary Fig. 1).

Several notable outliers were found from this data summary – all with lower-than-expected PT gene counts. These were *Laminaria digitata* (sublittoral brown alga), *Nitzschia alba* (coldwater diatom), and *Prototheca* spp. (parasitic algae). *N. alba* and *Prototheca* are both anomalous in the sense that they are apochlorotic members of their respective clades – having abandoned photosynthesis, their plastid genomes have become reduced. *L. digitata* does not have this feature. Probing this case further, we found that although its chloroplast genome is described as containing 139 coding sequences (CDSs) (Rana et al. 2019), its NCBI sequence NC_044689.1 only reports 107 CDSs. As its closest relatives in our dataset all have around 140 cpDNA CDSs, we marked this sequence as reflecting a possible missing data artefact, where not all genes are present in the reference sequence. The following analysis removes these outlier sequences, remembering that apochlorotic species depart from the trends we will report.

### Correlated departures from clade-wide gene count averages

To account for phylogenetic signal and pseudoreplication by relatedness (Paradis 2014; Maddison and FitzJohn 2015; Freckleton, Harvey, and Pagel 2002) in these observations, we changed variables to consider each species’ difference in oDNA gene count from its clade’s average, and performed phylogenetic linear modelling (PLM) accounting for relatedness. Thus, the mean MT and PT gene counts across a clade were computed, then subtracted from each species’ individual counts to obtain a difference (ΔMT and ΔPT respectively). A relationship between ΔMT and ΔPT, robust to phylogenetic correction, would suggest that those species with lower-than-expected MT gene counts also have lower-than-expected PT gene counts, and vice versa. In turn, this could suggest that similar pressures shape MT and PT gene counts at the species level.

A reasonably clear positive relationship (slope=0.16, p=0.0025) exists between ΔMT and ΔPT (Fig. 1B), although it must immediately be noted that a relatively small amount of the variance in one organelle gene count is explained by the other (R^2^ = 0.28). Supplementary Fig. 1 shows that, as expected, a linear mixed model with random effects assigned to clade shows a similar relationship.

To guard against artefacts from our chosen analysis method, we also analysed the raw MT and PT gene counts using PLM (without shifting by clade mean), and the relationship between ΔMT and ΔPT using a linear model (without phylogenetic correction via PLM). All variants of this process showed a positive relationship between mtDNA and ptDNA gene counts: slope 0.56, p=4.9×10^−5^ for PLM without clade mean shifting (equivalent to analysing the points in Fig. 1A accounting for relatedness); and slope 0.38, p=4.2×10^−15^ for a linear model after clade mean shifting (equivalent to equally weighting each point in Fig. 1B).

As Viridiplantae are the most sampled clade in our dataset, we also asked whether an MT-PT relationship was detectable within and without this clade. We used PLM to investigate the relationship between raw mtDNA and ptDNA gene counts in each case, finding significant relationships both within Viridiplantae (slope=0.15, p=0.0059) and without Viridiplantae (slope=2.0, p=6.5×10^−6^).

### Ecological and species-specific factors

Following work proposing ecological factors as shapers of oDNA evolution (García-Pascual, Nordbotten, and Johnston 2022; Giannakis, Richards, and Johnston 2023), we asked whether ecological factors further separated substructure in the MT-PT space (Fig. 2).

**Figure 2.**
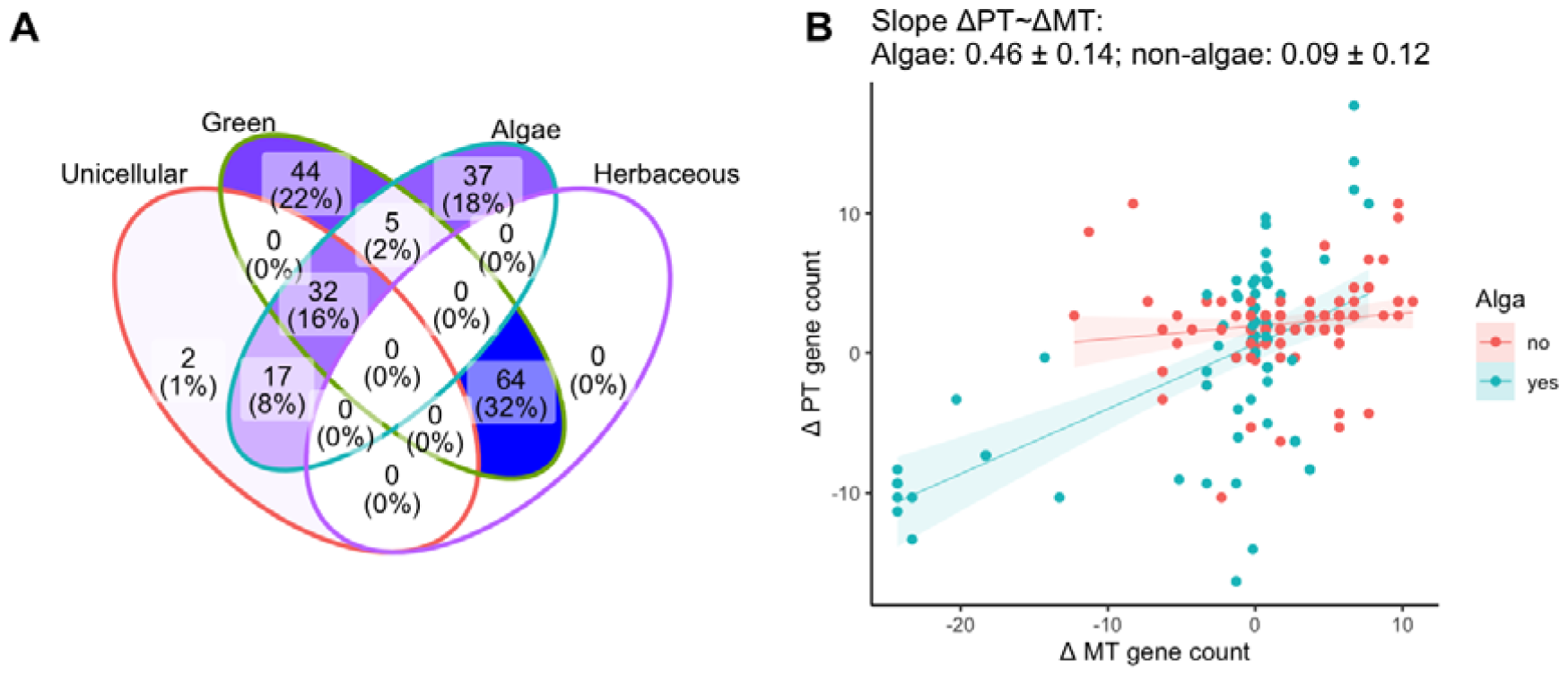
Ecological influences in ΔMT-ΔPT relationship. (A) Summary of the different classes of organism in the dataset. The two unicellular species that occupy no other sets are Apicomplexans. The herbaceous category is not applicable to algae or unicellular organisms. (B) ΔMT-ΔPT for organisms labelled as algae and organisms not labelled as algae. Statistically robust differences in MT-PT behaviour were not found for other arrangements of the features in (A).

Following findings in (Giannakis, Richards, and Johnston 2023), aligned with the features that most broadly distinguished entries in our dataset, we tested for differences between organisms with different characteristics: algae vs non-algae, unicellular vs multicellular organisms, green plants/algae vs others, herbaceous plants vs others, annual vs perennial lifestyle. For clade-corrected oDNA counts (ΔMT and ΔPT), a linear mixed model with random and gradients assigned to the algal category was slightly favoured by AIC over the fixed linear model (1154.7 vs 1157.5 for the fixed model). In other words, there is some statistical support for a picture where the relationship between ΔMT and ΔPT differs between algae and non-algae (Fig. 2B). We found no statistically robust differences in oDNA behaviour for other categorical differences between organisms.

### Gene-level relationships in mtDNA and ptDNA evolution

We next asked whether there was evidence for evolutionary relationships between specific MT and PT genes – for example, is there a particular PT gene that is always lost when a particular MT gene is also lost? To this end we used hierarchical clustering of the dataset to explore interdependence of PT-MT gene patterns (Supplementary Fig. 2). We also used Oncotree, an algorithm for inferring relationships in progressive evolutionary processes (Szabo and Boucher 2008), to explore evidence for gene-specific dependencies in our data. Oncotree is designed for and applied to features in cancer progression but fundamentally deals with progressive evolutionary changes in binary traits.

We found no evidence of gene-gene correlations or conditional dependencies via hierarchical clustering of the data (Fig. 3A), with MT and PT genes clustering separately across the retention profiles in our dataset, rather than clusters of MT and PT genes together which could indicate subset-specific correlations. Oncotree suggested some consistent ordering behaviour across organelles, where the loss of some MT genes (*ccmb, rps1, atp4, atp1, rps3, atp9*) is inferred to characteristically precede the loss of a collection of PT genes (Fig. 3B). This MT subset are not consistently among the most or least retained of organelle genes – their “retention indices” from (Giannakis et al. 2022) recording how many other genes are typically lost before them, take a range of intermediate values. The “dependent” PT subset overwhelmingly comprises highly-retained genes (Supp. Fig. 3). Most other genes from both organelles clustered separately in this analysis too, suggesting only limited gene-specific relationships between organelles.

**Figure 3.**
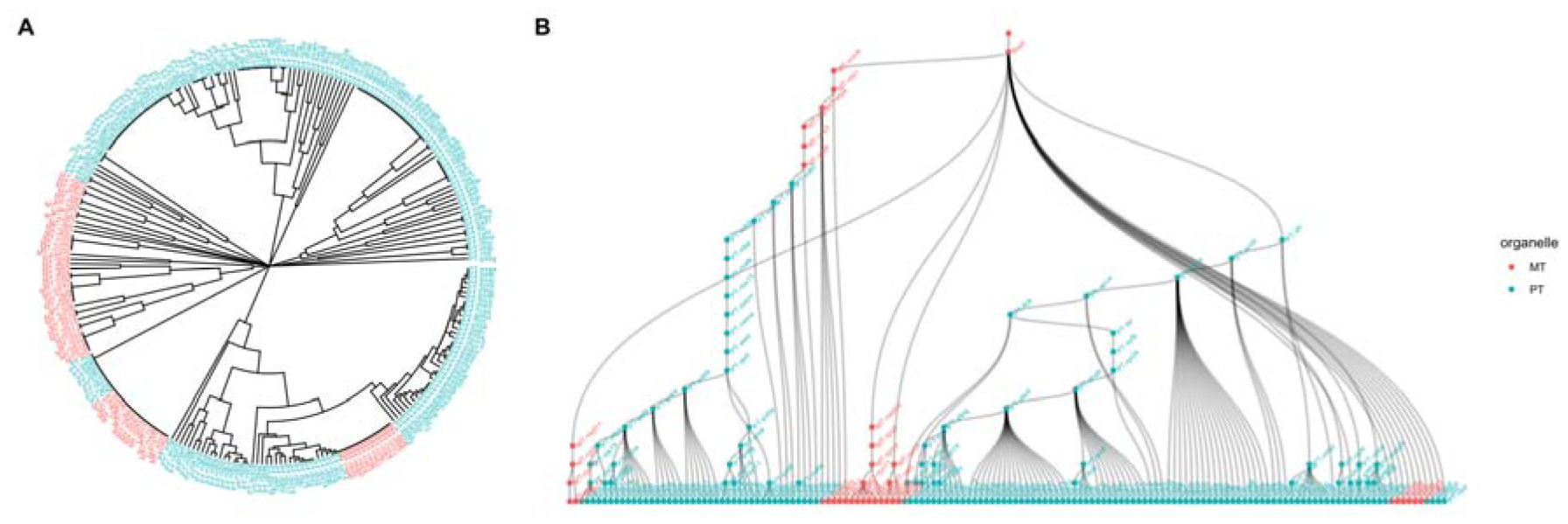
Limited evidence for gene-gene relationships across organelle types. (A) Hierarchical clustering of gene retention profiles in our dataset (Supplementary Fig. 2) produces a tree of relationships between genes that respects MT and PT differences. (B) Oncotree analysis of dependencies in MT-PT evolution, reporting inferred gene loss ordering in the dataset, shows mainly relationships within an organelle type (higher nodes are lost before connected lower nodes). Some MT genes (top, left of centre) precede a collection of PT genes, but the majority of genes cluster exclusively by organelle type.

## Discussion

Our work suggests that, controlling both for phylogenetic signal and the relatedness of individuals, there is some support for a picture where MT and PT gene counts are connected. This does not appear to be manifest through pronounced cross-organelle interactions between specific genes (where, for example, the loss of MT gene X always results in the loss of PT gene Y). Rather, it appears that the drivers of reductive evolution apply comparably, but largely independently, to the two organelles. These drivers appear to be connected to some aspects of organismal lifestyle – algae and non-algae, for example, display different MT-PT relationships.

This picture is compatible with recent results on both gene-specific (Giannakis et al. 2022; Johnston and Williams 2016) and species-specific (García-Pascual, Nordbotten, and Johnston 2022) drivers of oDNA genome evolution. Here, the same combination of features — hydrophobicity, GC content, and the centrality of a protein product in its functional complex — predict a “retention index” describing how readily a gene is lost in either mtDNA or ptDNA. In parallel, the demands placed on organelle control by a species’ environment determine how many oDNA genes it retains. Broadly, those with higher retention indices are retained even by species in less challenging environments; species in more challenging environments retain genes with progressively lower retention indices. There are, of course, many more determinants of the oDNA retention profile of individual species – from the outcomes of stochastic evolutionary processes to population genetic considerations, and doubtless other mechanisms which the above picture does not consider. Correspondingly, the gene-specific models in (Giannakis, Richards, and Johnston 2023; Johnston and Williams 2016) can explain just over half of the variance in gene retention indices (R^2^ = 0.51-0.60), and the relationship between MT and PT found here is certainly not clear-cut, with just over a quarter of variance in ΔPT explained by ΔMT (R^2^ = 0.28; Fig. 1B). This figure is of course after removing the apochlorotic and potentially misannotated outliers in our data, the inclusion of which would lead to still less variance being explained.

In correcting for systematic differences in gene count between deep-branching clades (Giannakis et al. 2022; Janouškovec et al. 2017) and additionally for the phylogenetic relationship between samples, we have attempted to align with the picture in (Maddison and FitzJohn 2015), of both identifying independent subsets of data and then accounting “internally” for phylogenetic signal. However, how best to account for dependencies across phylogenetically-embedded data is a topic of substantial discussion (Uyeda, Zenil-Ferguson, and Pennell 2018; Losos 2011); even well-principled comparative methods have shortcomings in the face of singular evolutionary events (Uyeda, Zenil-Ferguson, and Pennell 2018) and imperfect phylogenetic estimates (as we have here) and the distinction between pattern and process can challenge the application of these methods (Losos 2011) – discussed further in an oDNA context in (Giannakis, Richards, and Johnston 2023). In taking different combinations of clade-based and phylogenetic correction, and considering different subtrees of our data, we have attempted to be robust to particular choices of method. However, as more organelle genome data becomes available, testing alternative pictures like conditional dependencies between variables (Uyeda, Zenil-Ferguson, and Pennell 2018) will be an interesting line of inquiry.

The timescale of genome reduction must also be discussed. (Janouškovec et al. 2017), for example, give a compelling picture where the majority of mtDNA gene loss occurred early in the history of most eukaryotic clades, with modern-day mtDNA being more (in some cases almost completely) static. Despite evidence of ongoing movement of genetic information from oDNA to the nucleus (Timmis et al. 2004) – for example, nuclear mitochondrial sequences (NUMTs) from mtDNA in humans (Wei et al. 2022) and very frequent ptDNA transfer in plants (Stegemann et al. 2003; Bock and Timmis 2008) – the gene profiles of many eukaryotic clades seems quite fixed. The two pictures can be reconciled by noting that although NUMTs and NUPTs can enter the nuclear genome, the functionalization of these fragments into a working gene, and the subsequent removal of the organelle-encoded precursor, presumably involve many more genetic and regulatory steps and can reasonably be considered a rare event in many species – as in the limited transfer window hypothesis mentioned previously (Barbrook, Howe, and Purton 2006). Overall, the patterns in gene content observed across modern oDNA samples were presumably largely established in the rather distant evolutionary past. Hence, the processes we study here are historical; and while modern-day oDNA profiles are clearly compatible with modern-day environments, it was not modern-day environments that originally shaped them.

We have only considered protein-coding gene counts across species – a very coarse-grained structural feature of oDNA. Tremendous diversity also exists in oDNA length and proportion of coding content, genetic structure (gene orderings and repeat regions), and physical structure (linear, branching, circular, and fragmented molecules) of oDNA across eukaryotes (Roger, Muñoz-Gómez, and Kamikawa 2017; Keeling 2010). Explanations for this diversity include the mutational hazard hypothesis (Lynch, Koskella, and Schaack 2006; Smith 2016a), effects of oDNA recombination in some lineages (Edwards et al. 2021), and recent work suggesting a functional link between gene order and the control of gene expression in metazoa (Shtolz and Mishmar 2023); the interplay (or independence) of these mechanisms with the environmental influences we propose is an interesting target for future theoretical exploration.

Despite advances in sequencing protist oDNA, many clades are unrepresented in our dataset or only represented by a single species. Complete organelle records exist in NCBI for over 9k ptDNA sequences and over 2k mtDNA sequences – but the large majorities of these are from green plants and animals respectively (Smith 2016b). Organelle genome information from less represented clades – especially, for this study, photosynthetic protists – would be immensely valuable in verifying and refining these theories of organelle co-evolution. At the same time, such species play essential roles in marine and other ecosystems, and their photosynthetic and metabolic capacity and responses to environmental change are of profound importance on the warming and increasingly dynamic planet (Beardall and Raven 2004). More generally, oDNA features play an important role in the adaptability and evolvability of organisms in changing environments (Radzvilavicius and Johnston 2022; 2020), and environmental influences on photosynthesis and respiration are central to the biosphere’s response to climate change (Dusenge, Duarte, and Way 2019). Understanding these essential genomes further, and the advantages and disadvantages of gene retention in different environments, is highly desirable in the study of diverse eukaryotic responses to a changing world.

## Data and code availability

All data and code used in analysis and visualization is available at github.com/StochasticBiology/odna-coevolution/

## Acknowledgements

LR was supported by the BBSRC via the MIBTP Doctoral Training Scheme. This project has received funding from the European Research Council (ERC) under the European Union’s Horizon 2020 research and innovation programme (Grant agreement No. 805046 (EvoConBiO) to IGJ). KD was supported by the Trond Mohn Foundation [project HyperEvol under grant agreement No. TMS2021TMT09], through the Centre for Antimicrobial Resistance in Western Norway (CAMRIA) [TMS2020TMT11].

## Supplementary Information

**Supplementary Figure 1.**
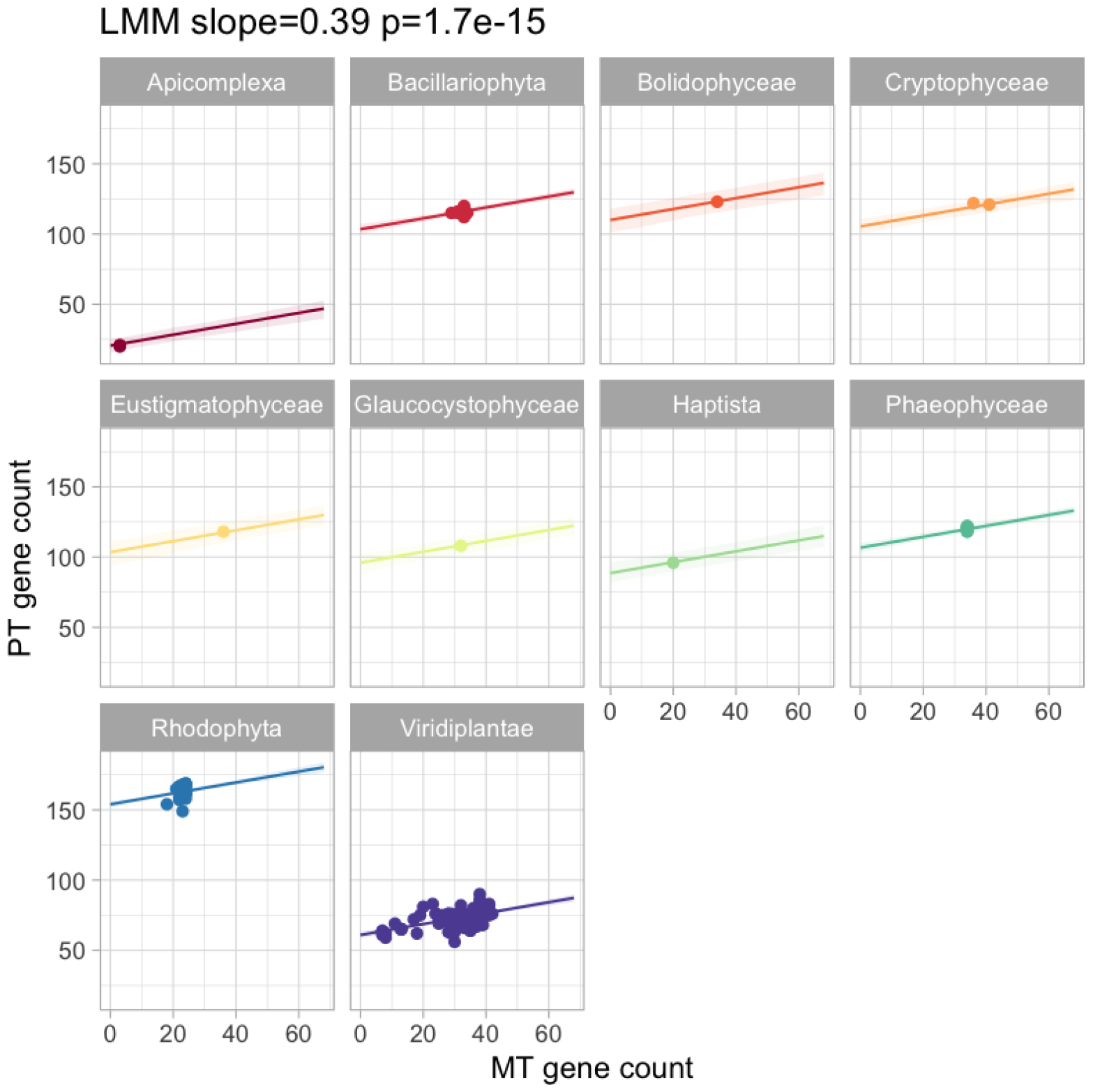
Linear mixed model for MT-PT relationship in raw gene counts, with random effects assigned to eukaryotic clade.

**Supplementary Figure 2.**
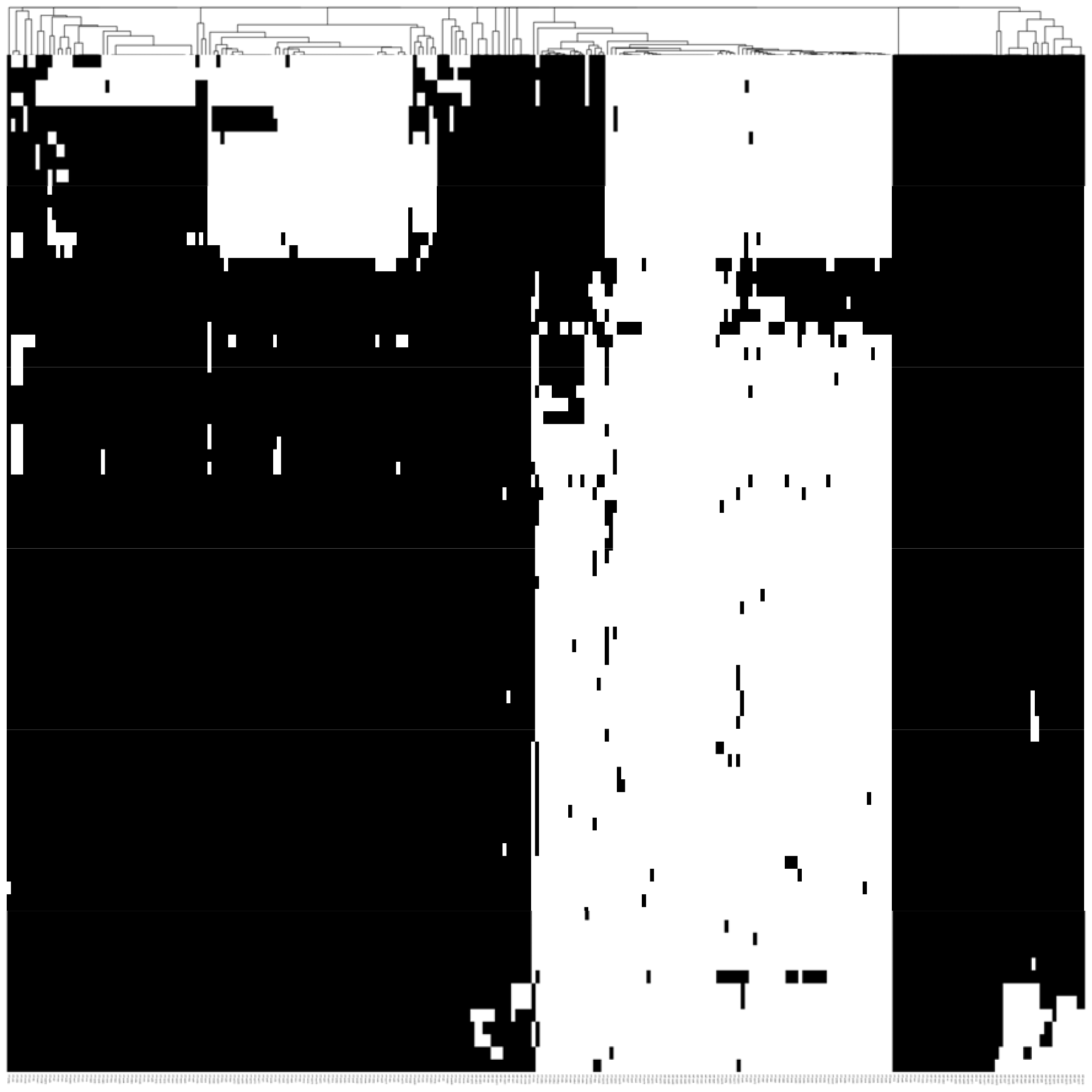
Heatmap from clustering in the combined MT-PT dataset; rows are retention profiles, columns are genes; black pixels are gene presence and white are gene absence. The dendrogram for gene clustering is reported in Fig. 3A.

**Supplementary Figure 3.**
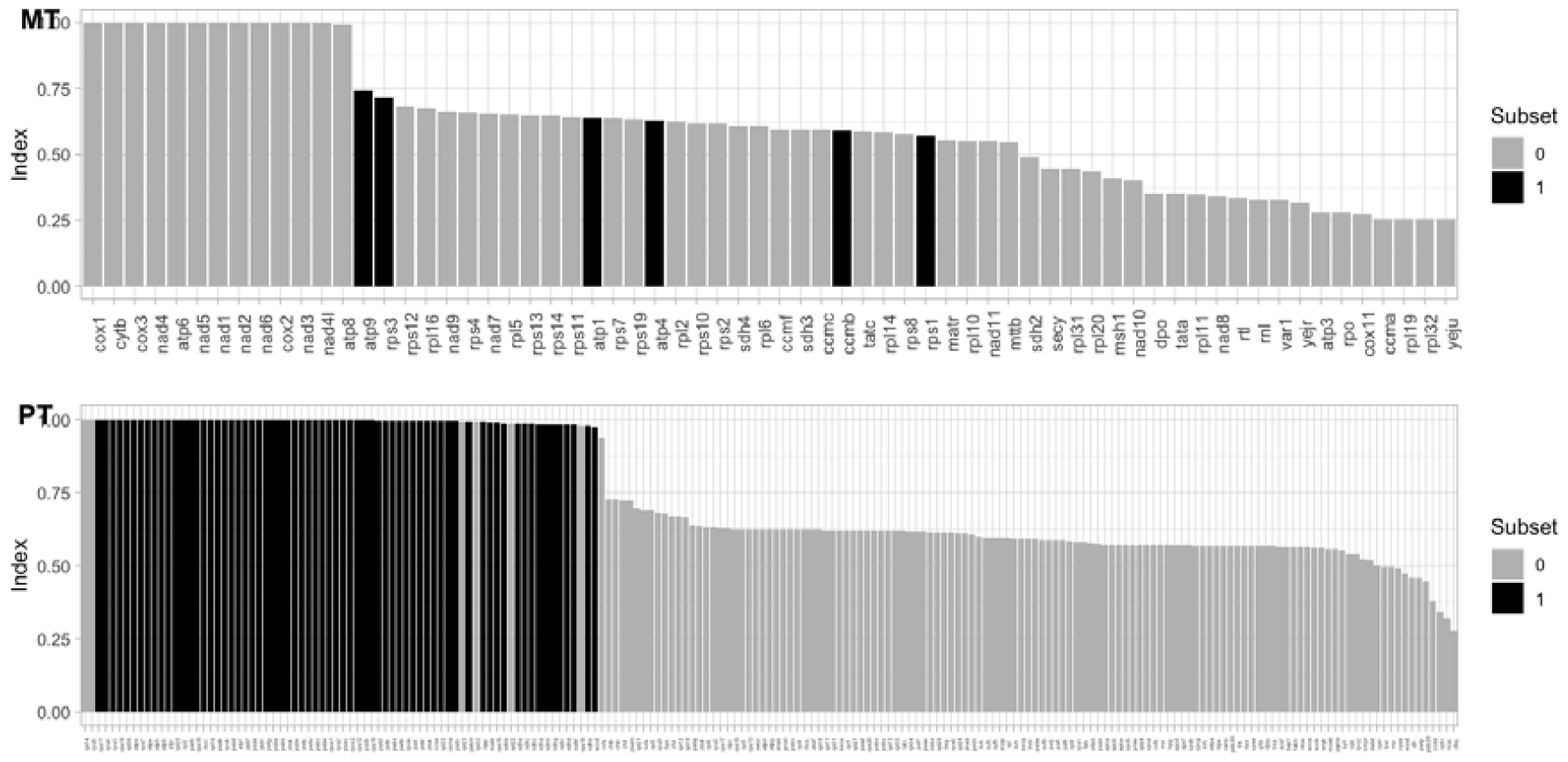
The subset of MT genes that Oncotree predicts to be lost before a subset of PT genes. Plots give the ranked “retention index” across oDNA genes from (Giannakis et al. 2022); genes with higher retention indices are inferred to be lost after genes with lower indices. The members of each subset identified by Oncotree are marked in black.

